# Myoelectric digit action decoding with multi-label, multi-class classification: an offline analysis

**DOI:** 10.1101/2020.03.24.005710

**Authors:** Agamemnon Krasoulis, Kianoush Nazarpour

## Abstract

The ultimate goal of machine learning-based myoelectric control is simultaneous and independent control of multiple degrees of freedom (DOFs), including wrist and digit artificial joints. For prosthetic finger control, regression-based methods are typically used to reconstruct position/velocity trajectories from surface electromyogram (EMG) signals. Although such methods have produced highly-accurate results in offline analyses, their success in real-time prosthesis control settings has been rather limited. In this work, we propose action decoding, a paradigm-shifting approach for independent, multi-digit movement intent decoding based on multi-label, multi-class classification. At each moment in time, our algorithm classifies movement action for each available DOF into one of three categories: open, close, or stall (i.e., no movement). Despite using a classifier as the decoder, arbitrary hand postures are possible with our approach. We analyse a public dataset previously recorded and published by us, comprising measurements from 10 able-bodied and two transradial amputee participants. We demonstrate the feasibility of using our proposed action decoding paradigm to predict movement action for all five digits as well as rotation of the thumb. We perform a systematic offline analysis by investigating the effect of various algorithmic parameters on decoding performance, such as feature selection and choice of classification algorithm and multi-output strategy. The outcomes of the offline analysis presented in this study will be used to inform the real-time implementation of our algorithm. In the future, we will further evaluate its efficacy with real-time control experiments involving upper-limb amputees.

---

Upper-limb loss can negatively impact an affected individual’s ability to perform activities of daily living. To mitigate this effect, prosthetic devices have historically aimed at restoring the appearance and basic functionality of a missing limb using artificial components. The advancement of robotics research in the last decades has led to the advent of upper-limb prostheses with highly-sophisticated mechanical capabilities. Sensing technologies and control algorithms, however, have not kept pace; as a result, they currently impose a bottleneck on the control dexterity enjoyed by prosthesis users.

Prosthetic hands are typically controlled using muscle activity signals, called electromyograms (EMGs), recorded on the skin surface. Traditionally, clinical solutions have deployed simple, amplitude-based control algorithms which rely on monitoring the activity of a pair of antagonist remnant muscles. The amplitude of each recorded muscle is usually linked to the activation of a specific function, for example, wrist rotation or hand opening/closing. To access a different function, the user has to switch between the available modes by using a trigger signal, such as muscle co-contraction^1^. Although this algorithm has proven robust, it results in limited control and can also be non-intuitive and cumbersome for the end-user. Unfortunately, this leads to an increased rate of prosthesis rejections^2^.

To improve control dexterity, machine learning algorithms can be used to decipher movement intent from EMG recordings. Typically, a classifier is used to map features extracted from multiple EMG channels onto a discrete output variable encoding grasp type and/or other prosthesis functions. This paradigm has been highly-successful and, in the last decade, has found its way towards commercial adoption^3^. One caveat of this approach is that it can only support a single function activation at a time. That is, to achieve outcomes that require activating more than one prosthesis functions, for example, wrist rotation and hand closing. the user has to trigger a sequence of commands. This results in sub-optimal and unnatural control. To address this issue, simultaneous (i.e., *multi-label*) classification has been proposed as a means of decoding multiple hand and wrist functions together, hence resulting in greater flexibility and dexterity^4–9^.

Intuitive selection and activation of grasp and wrist functions can be greatly beneficial for the user in performing activities of daily living. Yet, this paradigm results in severe prosthesis under-actuation and can thus offer much less functionality and dexterity than a natural hand. From a technical perspective, the ultimate goal of the myoelectric control field is to approximate this dexterity via simultaneous and independent control of multiple degrees of freedom (DOFs) in a continuous space. To that end, several groups, including us, have used regression-based methods to reconstruct wrist kinematic trajectories^10–14^, finger positions^15–21^ and velocities^22, 23^, as well as fingertip forces^24–27^. Only a few studies have, however, thus far demonstrated the feasibility of real-time prosthetic finger control in amputee users^16, 20, 21^. This indicates that independent prosthetic digit control is, indeed, a challenging problem, which calls for new and more efficient methods of tackling it.

In this work, we propose a novel approach for simultaneous and independent control of prosthetic digits. Our algorithm is based on multi-label, multi-class classification. This is in contrast with previous work in this area, which has focused on the use of multi-output regression algorithms to achieve the same goal. At each time step, our algorithm uses surface EMG features to decode movement intent for each available DOF into one of three classes: open, close, or stall (i.e., no movement). Our motivation is to replace the multi-output regressor with a multi-label classifier with the aim of simplifying the decoding part and improve control performance. We term our approach *action control*, since it is based on decoding digit actions rather than positions and/or velocities. A schematic of the approach is shown in Figure 1.

**Figure 1.**
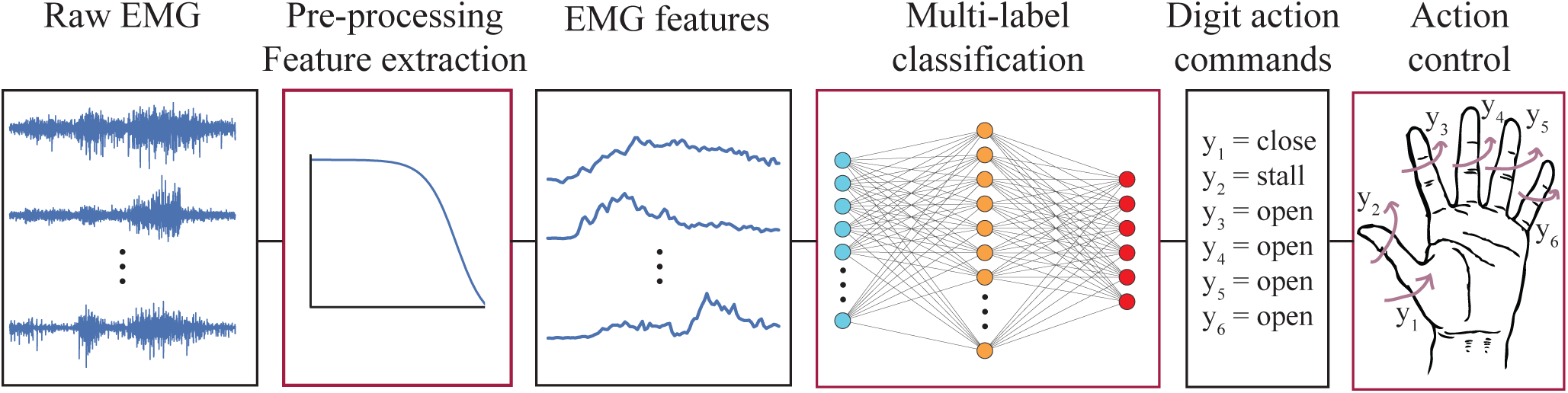
Action decoding paradigm. Multi-channel raw EMG measurements are pre-processed and fed as inputs into a multi-label classifier. The classifier has six outputs corresponding to the following DOFs: thumb rotation and flexion/extension of the thumb, index, middle, ring, and little digits. For each DOF, the algorithm classifies movement intent into one of three actions: open, close, or stall (i.e., no movement). Predictions can then be used to control the digits of a prosthesis using discrete actions.

We have previously evaluated the proposed action controller in a robotic hand tele-operation task with a data glove and found that it can achieve comparable performance to digit position (i.e., joint angle) control^28^. Here, we provide a first implementation of the method in the context of myoelectric decoding. We demonstrate the feasibility of using surface EMG measurements to decode digit actions. We also perform a systematic offline analysis investigating several aspects of the method, including feature selection and choice of classifier and multi-output strategy. With regard to the latter, we evaluate the efficacy of a state-of-the-art method for multi-label classification, namely, classifier chains (CCs), which takes output dependencies into account when making predictions. The outcomes of our analysis are used to inform the real-time implementation of the algorithm, which we present in a separate study^29^.

## Results

We analysed data from 10 able-bodied and two transradial amputee subjects. We extracted features from 16 surface EMG channels and used them to decode digit movement (i.e., actions) for the following DOFs: thumb rotation and flexion/extension of the thumb, index, middle, ring, and little digits.

As a first step, we performed an exhaustive feature analysis including a large number of commonly used time-domain EMG features. For each subject, features were ranked using a forward feature selection algorithm and we computed mean ranks for all features. The results of this analysis are presented in Figure 2(a). The highest-performing feature was Wilson amplitude (WAmp), followed by log-variance (LogVar) and Hjorth (Hjorth) parameters. Figure 2(b) shows average classification performance by means of F1-score for an increasing number of added features. We observed a plateau in performance after including six to eight features. Based on this observation, we selected the seven most highly-ranked features for the rest of our). analysis: WAmp, LogVar, Hjorth, kurtosis (Kurt), auto-regressive (AR) coefficients, waveform length (WL), and skewness (Skew).

**Figure 2.**
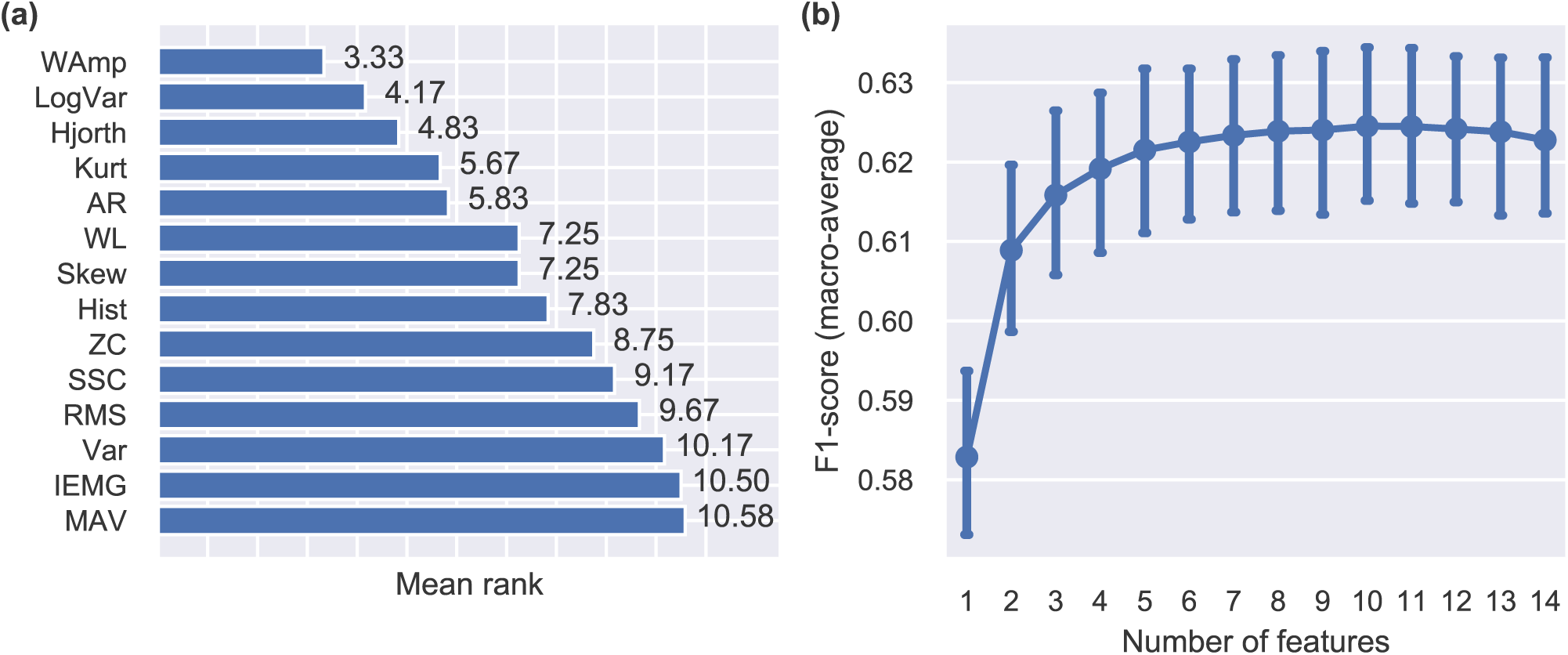
Feature analysis. **(a)** Average ranking for individual features using the sequential forward selection method. The procedure was run independently for each participant and average rankings were computed (*n* = 12). **(b)** Performance as a function of the number of features used for decoding. Higher F1-scores indicate better performance. Points, means; error bars, standard errors estimated via bootstrapping (1000 iterations).

Next, we investigated the potential of using CCs and ensembles of CCs to improve classification performance by exploiting label (i.e., output) dependencies. The results of this analysis are presented in Figure 3(a). For each participant, we report the performance of the best- and worst-performing CCs, as well as average performance of ensemble CCs consisting of 10 chains with random label orders. We compare the performance of CCs end ensemble CCs to that of multiple independent classifiers. In all cases, we used linear discriminant analysis (LDA) models as the base classifiers for the CCs. We use F1-score (see Methods section) throughout as our preferred performance measure, unless noted otherwise.

**Figure 3.**
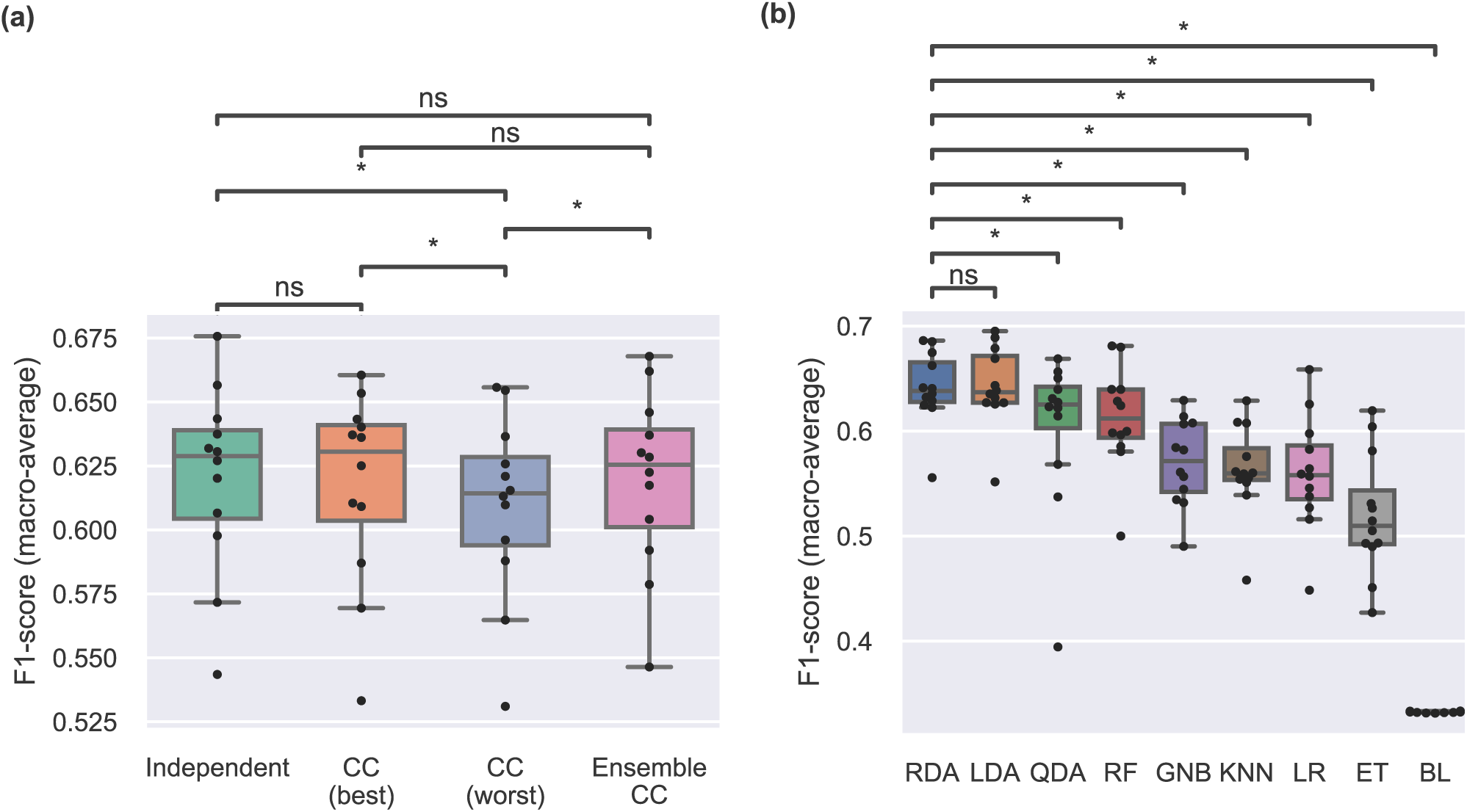
Performance comparisons. **(a)** Comparison of multi-output classification strategies using an LDA classifier. CC best and CC worst correspond to best- and worst-performing single CC models, respectively. For each participant, 100 random chains were generated. Extended CC corresponds to a model with 10 random chains. Straight lines, medians; solid boxes, interquartile ranges; whiskers, overall ranges of non-outlier data; dots, individual data points (*n* = 12); asterisk, *p* < 0.05; n.s., *p* > 0.05. **(b)** Comparison of classification algorithms using F1-score and the independent multi-output strategy. Classifiers are presented in order of decreasing median performance and statistical comparisons are performed only against the highest-performing algorithm. CC, classifier chain; RDA, regularised discriminant analysis; LDA, linear discriminant analysis; QDA, quadratic discriminant analysis; RF, random forest; GNB, Gaussian naive Bayes; KNN, k-nearest neighbours; LR, logistic regression; ET, extra trees; BL, baseline.

We did not observe a difference in F1-score between the CC-best and independent output classifiers (*p* = 0.51). Moreover, performance was not statistically different when we used ensembles of 10 CCs (*p* = 0.61). F1-scores for CC-worst were statistically lower than independent (*p* = 0.02), CC-best (*p* = 0.03), and ensemble CCs (*p* = 0.02). This finding indicates that a poor label ordering can in fact decrease performance, as compared to independent classifiers. Additional results from this analysis using other performance metrics are provided in Supplementary Figure S1.

Figure 3(b) summarises the results of the classifier benchmark analysis using independent output classifiers. The highest median performance was achieved by regularised discriminant analysis (RDA) (*F*1_*med*_ = 0.64), closely followed by LDA. We performed statistical comparisons between the highest-performing algorithm (i.e., RDA) and all other classifiers and found that RDA significantly outperformed all classifiers except for LDA. All classifiers performed higher than chance.

Figure 4 shows average confusion matrices obtained with the independent output RDA classifier for individual DOFs. The best performance was achieved for the ring digit, followed by the middle, index, and thumb digits. The lowest performance was achieved for the thumb rotation and little digit flexion/extension DOFs.

**Figure 4.**
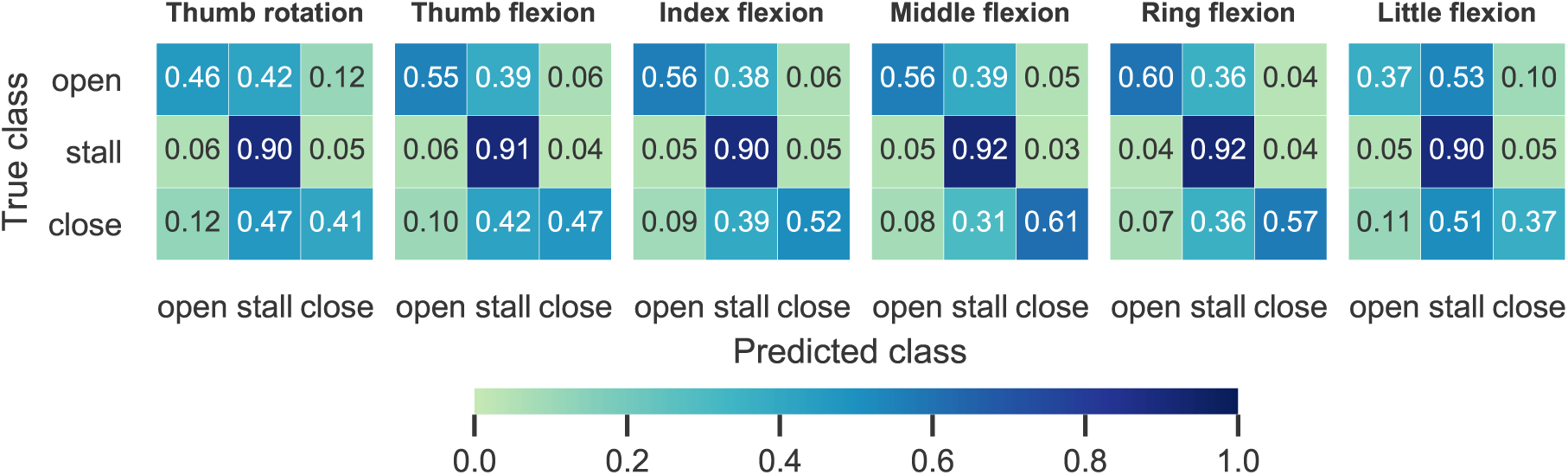
Confusion matrices. For each DOF, the average confusion matrices obtained with the multi-output independent RDA classifier are shown. Colour bar and annotated scores indicate normalised prediction rates.

Additional results from the benchmark analysis are provided in the supplementary material: Figure S2 shows algorithmic comparisons for both independent classifiers and CCs including additional performance metrics; Figure S3 shows performance of the two highest-performing algorithms (e.g., LDA and RDA) for individual subjects; and Figure S4 summarises performance of all classifiers for individual DOFs using independent classifiers for each output.

## Discussion

In this work, we introduced a novel paradigm for upper-limb prosthetic digit control using surface EMG signals. In the proposed approach, EMG features are used to decode discrete digit actions via multi-label classification. At each time step, the algorithm classifies movement intent for each DOF into one of three categories: open, close, or stall (i.e., no movement). We have previously evaluated this type of controller in a robotic hand tele-operation task with a data glove and have found that it can achieve comparable performance to digit position control^28^. The aim of this study was twofold: 1) to demonstrate the feasibility of using surface EMG signals from the forearm to decode digit actions; and 2) to carry-out a systematic offline investigation prior to implementing the algorithm in real-time and testing it with upper-limb amputees.

We have shown that it is feasible, in principle, to decode digit actions from surface EMG signals. The median F1-score of the best-performing configuration (i.e., independent multi-output RDA classifiers) was *F*1_*med*_ = 0.64. The median Hamming score, exact match ratio, and macro-average precision and recall scores were all significantly and substantially higher than chance (supplementary material). We observed the highest performance for the ring and middle digits (Figure 4). This is in agreement with previous work on regression-based reconstruction of digit position/velocity trajectories^17, 20, 22^. This finding is expected from a physiological perspective, given that these two digits are controlled by extrinsic superficial muscles, as opposed to the thumb, for example, which is controlled by intrinsic and deep extrinsic muscles, both of which are not easily accessible from the surface of the forearm.

Our offline investigation scrutinised several important aspects of the method, including feature analysis, choice of decoding algorithm, and evaluation of two different multi-output strategies. It has been previously demonstrated that both feature selection and choice of classifier can substantially influence the performance of myoelectric classification systems^30, 31^. In line with previous reports, which were mainly concerned with upper-limb motion/grip classification^8, 30–32^, we found that discriminant analysis-based classifiers, such as LDA and RDA, can achieve the highest level of performance. The results of our feature selection/ranking analysis (Figure 2(a)) are also largely in agreement with previous reports from the motion classification literature^31, 33^.

In multi-label classification settings, it is often desirable to exploit output dependencies to improve decoding performance. In this regard, we investigated the potential of using the state-of-the-art method of CCs to improve classification. The exact match ratio was marginally improved with both CCs and extended CCs (see supplementary material). Nevertheless, we did not observe an increase in F1-score using either method. The improvement in exact match ratio is expected, since it has been theoretically shown that CCs maximise this metric exactly by equivalently minimising 0/1 loss^34^. On the other hand, when labels are evaluated independently, as with macro-average F1-score, there is no guarantee that CCs will outperform independent classifiers, although this may often happen in practice^35^. We attribute the ineffectiveness of CCs in our case to the fact that the dataset comprised both single-digit exercises as well as full-hand grips (five exercises of each type). Including a large number of single-digit motions results in outputs becoming largely independent and, thus, there is less structure in the output domain that CCs can exploit to leverage performance.

In myoelectric control, multi-label classification has been previously used to decode simultaneous wrist and hand motions^4–9^. For control of prosthetic digits, however, previous efforts have focused on using multi-output regression to reconstruct position^15–18, 20^, velocity^22, 23^ or fingertip force trajectories^25–27^. One study has previously adopted a similar approach to ours^36^, but with three main differences: firstly, the labels corresponded to digit positions instead of actions; secondly, labels were binary (i.e., digits could be fully open or closed), whereas with our approach actions can take three values (i.e., open, close, or stall); finally, due to using binary outputs, a controller using the approach proposed previously would be limited to extreme digit positions. In contrast, with our action-based decoder, intermediate positions are possible. We have previously shown in a hand tele-operation task with a data glove^28^ that by using a small action step (i.e., digit position increase/decrease step), control becomes approximately continuous. In other words, despite using a classifier as the decoder, our approach allows for arbitrary hand configurations. From a control theory perspective, our action-based paradigm can be viewed as an extreme discretised case of velocity decoding; velocity can either be zero, or take a constant value, which is only parametrised by its sign.

Our study has two limitations: firstly, it was limited to offline analyses. It is well-accepted in the myoelectric control community that offline performance measures are not always a good proxy of real-time control performance^9, 13, 20, 37, 38^. Therefore, it is imperative to evaluate control algorithms with real-time implementations and user-in-the-loop experiments. Given that we have introduced a completely novel paradigm for prosthetic digit control, the main purpose of this work was to systematically explore different parameters of the method and lay the groundwork for the subsequent real-time implementation. We report our implementation and evaluation of the proposed algorithm with amputee participants in a separate study^29^.

The second limitation of the study was that we did not consider neural networks in our classifier benchmarks. Neural networks can naturally handle multi-label, multi-class classification problems via appropriate design of the output layers. For our application, it is likely that parameter sharing in the early layers of a network may benefit overall performance by optimising a combination of ouput loss functions, one for each DOF (i.e., multi-task learning). This is currently seen as a future research direction.

In conclusion, we have proposed a new paradigm for prosthetic digit control based on multi-label, multi-class classification. We have demonstrated the feasibility of decoding actions for all five digits and rotation of the thumb using surface EMG measurements recorded on the forearm in both able-bodied and transradial amputee participants. Our algorithm warrants further investigation with real-time, user-in-the-loop experiments with upper-limb amputees.

## Methods

### Dataset

The dataset used in the study was previously collected and made publicly available by us^20^. For completeness, we briefly describe here the experimental protocol and refer the reader to the original study for more details.

Ten able-bodied and two right-arm transradial (i.e., below-elbow) amputee participants were included in the study. All able-bodied participants were right-hand dominant. All participants read an information sheet and gave written consent prior to the experimental sessions.

We placed 16 Delsys Trigno surface EMG sensors (Delsys, Inc.) on the participants’ skin below the right elbow arranged in two rows of eight equidistant electrodes and without targeting specific muscles. Prior to electrode placement, we cleansed participants’ skin using 70% isopropyl alcohol. We used adhesive elastic bandage to secure the locations of the electrodes throughout the sessions. The sampling rate of EMG data acquisition was fixed at 1111 Hz.

In addition, we recorded hand kinematic data using an 18-DOF Cyberglove II data glove (CyberGlove Systems, LLC), which we placed on the participants’ left hand. Data glove measurements were calibrated for each participant using a quick calibration routine provided by the manufacturer. The sampling rate of glove data acquisition was fixed at 25 Hz.

Participants sat comfortably on an office chair and rested both arms on a table. They were asked to perform a series of bilateral mirrored hand exercises whilst these were instructed on a computer display placed approximately 1 m in front of them. The following single-finger and full-hand exercises were included: thumb abduction/adduction; thumb, index, middle, and combined ring and little fingers flexion/extension; cylindrical, lateral, and tripod grips; and index pointer. Three blocks of exercises were recorded for each participant: datasets A and B comprised 10 repetitions of each exercise and dataset C only two. Consecutive trials were interleaved with 3 s of rest. The experimental protocol and apparatus used are shown in Figure 5.

**Figure 5.**
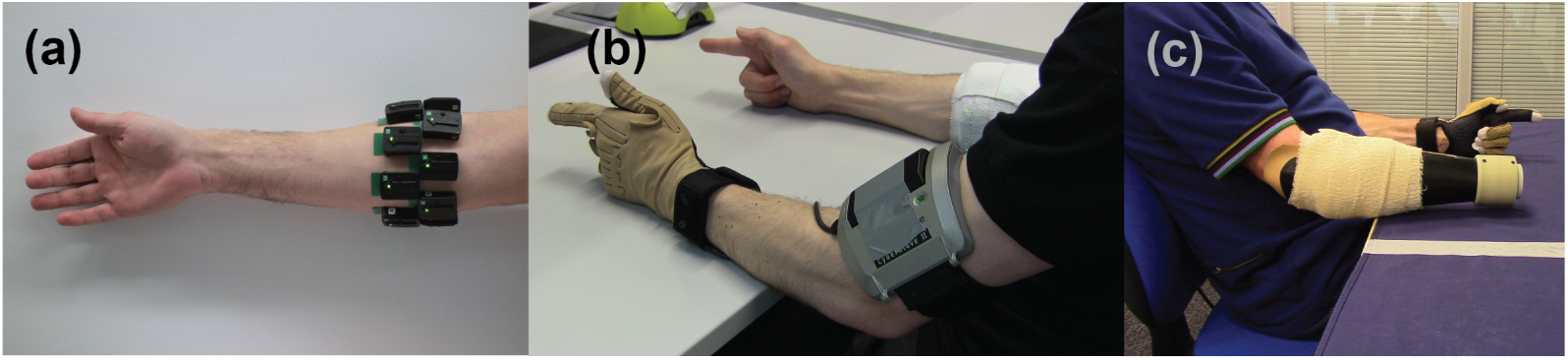
Data collection. **(a)** Sixteen wireless EMG sensors were placed on the surface of the skin and were arranged in two rows of eight equidistant sensors below the elbow. **(b)**-**(c)** Hand kinematic data were collected using an instrumented data glove placed on the contralateral side. Participants were instructed to perform bilateral mirrored movements.

### Pre-processing

Myoelectric and glove data were upsampled to 2 kHz and synchronised using linear interpolation. We processed myoelectric data using an overlapping window approach. We set the window length to 128 ms and the increment to 50 ms (i.e., approximately 60% overlap). We filtered the EMG signals using a 4^th^-order band-pass Butterworth digital filter with lower and upper cutoff frequencies of 10 and 500 Hz, respectively.

We transformed the calibrated glove measurements into joint angles for the following six DOFs using a linear mapping: thumb rotation, flexion/extension of the thumb, index, middle, ring, and little digits. Joint angles were then normalised in the range 0 to 1, with 0 corresponding to a DOF being fully open (or thumb rotator fully reposed) and 1 to fully closed (thumb rotator fully opposed). Finally, we smoothed the calibrated glove measurements with a 1^st^-order low-pass Butterworth filter with cutoff frequency of 1 Hz.

### Digit action estimation from joint angle trajectories

To extract digit action labels from calibrated glove data we used the following procedure. Firstly, we estimated joint velocities by computing the first-order difference of normalised joint angle trajectories. We then thresholded the computed differences using a tolerance *ε* = 0.004, such that joint velocities larger than *ε* were assigned the “close” label and velocities smaller than *−ε* were assigned the “open” label. Velocity values in the range [*− ε, ε*] were assigned the “stall” label, corresponding to no movement. Finally, for joint angles less than 7.5% away from either boundary (0 or 1), we assumed that the respective actions were “open” and “close”, respectively, regardless of the joint velocity. The digit action estimation procedure was performed independently for each DOF. An illustration is provided in Figure 6 using data from one participant as an example.

**Figure 6.**
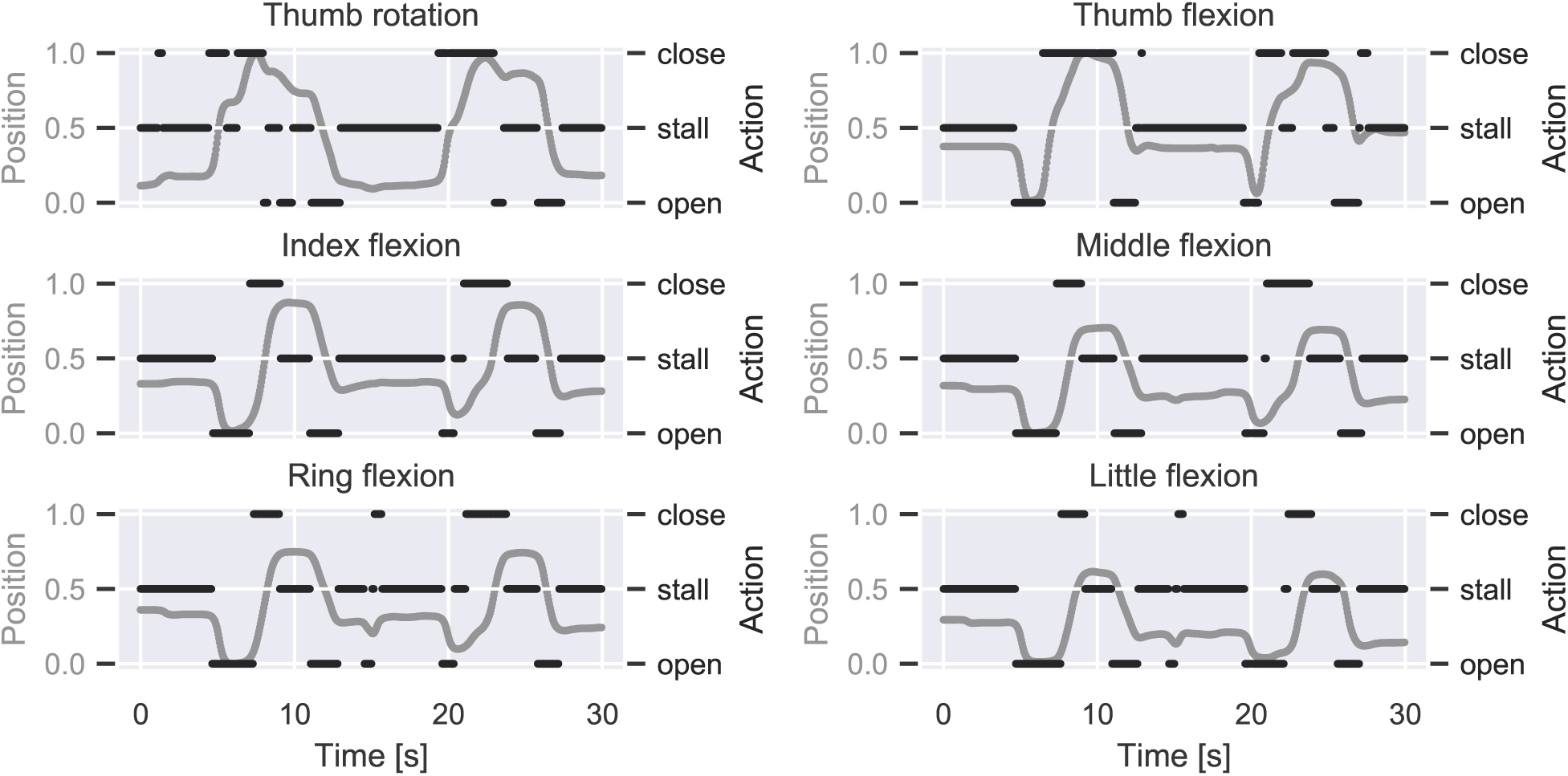
Finger action estimation from glove data. The mapping from position to action space is demonstrated using data from one participant. The position trajectory for each DOF (left y-axes) is normalised between 0 (i.e., fully open) and 1 (i.e., fully closed). The first-order discrete position difference is computed and transformed into action via thresholding (right y-axes). The shown excerpt corresponds to two repetitions of the cylindrical grasp exercise.

### Performance evaluation and metrics

We considered a range of performance metrics to characterise classification performance. In a multi-label classification setting, the following metrics are commonly used^39–41^: 1) exact match ratio or accuracy is the percentage of samples that have all their labels correctly classified; 2) hamming score is the fraction of correctly classified labels to the total number of labels; 3)precision, recall and F1-score (i.e., harmonic mean of precision and recall) can be used in a similarly way to multi-class classification, that is, either on a per-label basis or by using an appropriate method to average across labels. Macro- and micro-averaging are common choices; macro-averaging computes the metric of interest for each label independently and averages across labels, whereas micro-averaging aggregates contributions from each label to compute the average metric.

The exact match ratio is a strict measure, since it requires all labels to be correctly classified for a sample to be considered correct. Given that the number of labels in our case was relatively large (i.e., six), we did not consider this measure as our main evaluation metric. On the other hand, training/testing datasets for individual participants were highly-imbalanced, due to domination of the “stall” class over the “open” and “close” classes. Therefore, hamming loss was not an appropriate evaluation metric either. Taking the above into consideration, we selected F1-score as our main performance measure and used macro-averaging to account both for multiple labels, as well as multiple classes within each label. We report macro-average F1-scores in the main part of our analysis; additional results are reported in the supplementary material using the following metrics: exact match ratio (i.e., accuracy), hamming score, recall (macro-average) and precision (macro-average).

### EMG feature extraction and selection

We experimented with a large group of time-domain EMG features. In the feature analysis investigation, we included the following time-domain features: mean absolute value (MAV)^42^, waveform length (WL)^31^, rate of zero-crossings (ZC)^43^, slope sign changes (SSC)^31^, Wilson amplitude (WAmp)^43^, root mean square (RMS)^31^, integrated EMG (IEMG)^43^, variance (Var)^43^, log-variance (LogVar)^11^, kurtosis (Kurt)^44^, skewness (Skew)^45^, 4^th^-order auto-regressive (AR) coefficients^43^, histogram (Hist) counts^43^ (*n*_*bins*_ = 5), and Hjorth (Hjorth) parameters (i.e., activity, mobility, and complexity)^46^.

For the feature analysis investigation, we used a modified version of the sequential forward feature selection algorithm^47^. The algorithm was initialised with an empty feature set. Within each iteration, all available features were added to the pool, one at a time, and the respective F1-scores were estimated. The feature that yielded the highest performance was added to the pool and the procedure was repeated until all features were included. In that way, each EMG feature was assigned a rank, which was equal to the order that it was added to the pool. The forward selection procedure was performed independently for each participant and respective feature ranks were averaged. For each participant, models were fitted using independent multi-output LDA classifiers on dataset A and were evaluated on dataset B.

Based upon the results of the feature selection analysis (Figure 2), we used the following features in the rest of the study: WAmp, LogVar, Hjorth, Kurt, AR, WL, and Skew.

### Classifier chain analysis

Classifier chains is a popular machine learning tool for multi-label classification that takes into account label dependencies. We briefly describe the method here and refer the interested reader to the original paper for more details^35^.

Given a set of labels ℒ, a classifier chain model learns |ℒ | classifiers, which are linked in a chain. Firstly, the label chain (i.e., order of labels) needs to be defined: {*L*_1_, *L*_2_, …, *L*_|ℒ |_}. The first classifier in the chain *L*_1_ is then fitted using input features only. The ground truth data from *L*_1_ are then included as an additional input feature for training the second classifier in the chain *L*_2_. This process is repeated for all remaining labels in the chain by including for each label *L*_*n*_ ground truth data from previous labels in the chain {*L*_1_, *L*_2_, … *L*_*n−*1_}. For inference, the same procedure is followed, except predictions from previous labels in the chain are used at each step. A popular variant of the method is the ensemble of classifier chains, whereby several classifier chain models are trained with random orders of labels and their predictions are aggregated using a voting scheme.

In our application, the number of labels (i.e., outputs) was |ℒ| = 6, that is, the number of DOFs: thumb opposition/reposition and flexion/extension of thumb, index, middle, ring, and little digits. For each label, the set of classes was *ℐ* = {*open, close, stall*} and thus the number of classes was |*ℐ*| = 3. In our classifier chain analysis, we tested all possible orders of labels, that is, a total of 6! = 720 chains. For each participant, we report performance for the best- and worst-performing chains in terms of F1-score. In addition, we report best performance from a set of 100 ensemble classifier chain models, each trained with a random set of 10 label orders. We implemented the ensemble classifier chains using a *soft* voting scheme, which predicts class labels based on the predicted probabilities from each chain in the ensemble. We compared the performance of CCs and ensemble CCs to that of independent multi-label classification, whereby an independent classifier is trained for each output. This method is often referred to as *binary relevance*^39, 40^ when dealing with multi-label binary classification problems. Note, however, that this term is not appropriate for our problem, which is multi-label and multi-class. Therefore, we refer to this strategy as independent multi-output classification.

### Classifier training and hyper-parameter optimisation

We considered a wide range of classifiers in our classification benchmark analysis. With few exceptions (e.g., LDA, quadratic discriminant analysis (QDA) and Gaussian naive Bayes (GNB)), most algorithms have hyper-parameters which were systematically tuned with hold-out validation. For each participant, we fitted models on dataset A and tuned hyper-parameters using randomised search with 50 iterations on dataset B. We finally report performance on dataset C. We performed 10 independent runs for each participant/classifier experiment and report average performance results across runs. Baseline performance was assessed using a dummy classifier always predicting the “stall” class for each label, which was the dominating class in the training dataset. The list of algorithms used in the benchmark along with hyper-parameters and respective search ranges for each classifier are provided in Table S1.

Using 16 EMG electrodes and the optimal feature set identified as part of the feature selection analysis (i.e., WAmp, LogVar, Hjorth, Kurt, 4^th^-order AR coefficients, WL, and Skew), the input dimensionality was *D* = 192. For the k-nearest neighbours (KNN) and logistic regression (LR) classifiers we reduced the input dimensionality to *D* = 50 using principal component analysis to speed-up training. The approximate number of samples for the training and validation sets was 22,000 and for the test set was 5,000. We performed model training and testing in `Python` using the `scikit-learn` library^48^ and custom-written code.

## Statistical analysis

We used two-sided Wilcoxon signed-rank tests to compare performance between pairs of classifiers. All comparisons were performed at the population level using participant-average scores, thus the number of samples was *n* = 12. To account for multiple comparisons, we used the Holm-Bonferroni correction method. Statistical analysis was performed in in `Python` using the `Pingouin` library^49^.

## Data availability

The dataset used in the study is available as part of the NINAPRO database (http://ninapro.hevs.ch/DB8) and the Newcastle University data repository (https://doi.org/10.25405/data.ncl.9577598.v1). Meta-data are available from the authors upon reasonable request.

## Acknowledgements

AK and KN are supported by the Engineering and Physical Sciences Research Council (EPSRC) under grant EP/R004242/1.

## Author contributions statement

AK designed the study. AK analysed the data and prepared figures. AK wrote the manuscript. AK and KN read and approved the final manuscript.

## Additional information

### Supplementary information

accompanies this paper.

