## Supplementary information for "Myoelectric digit action decoding with multi-label, multi-class classification: an offline analysis"

### Supplementary figures

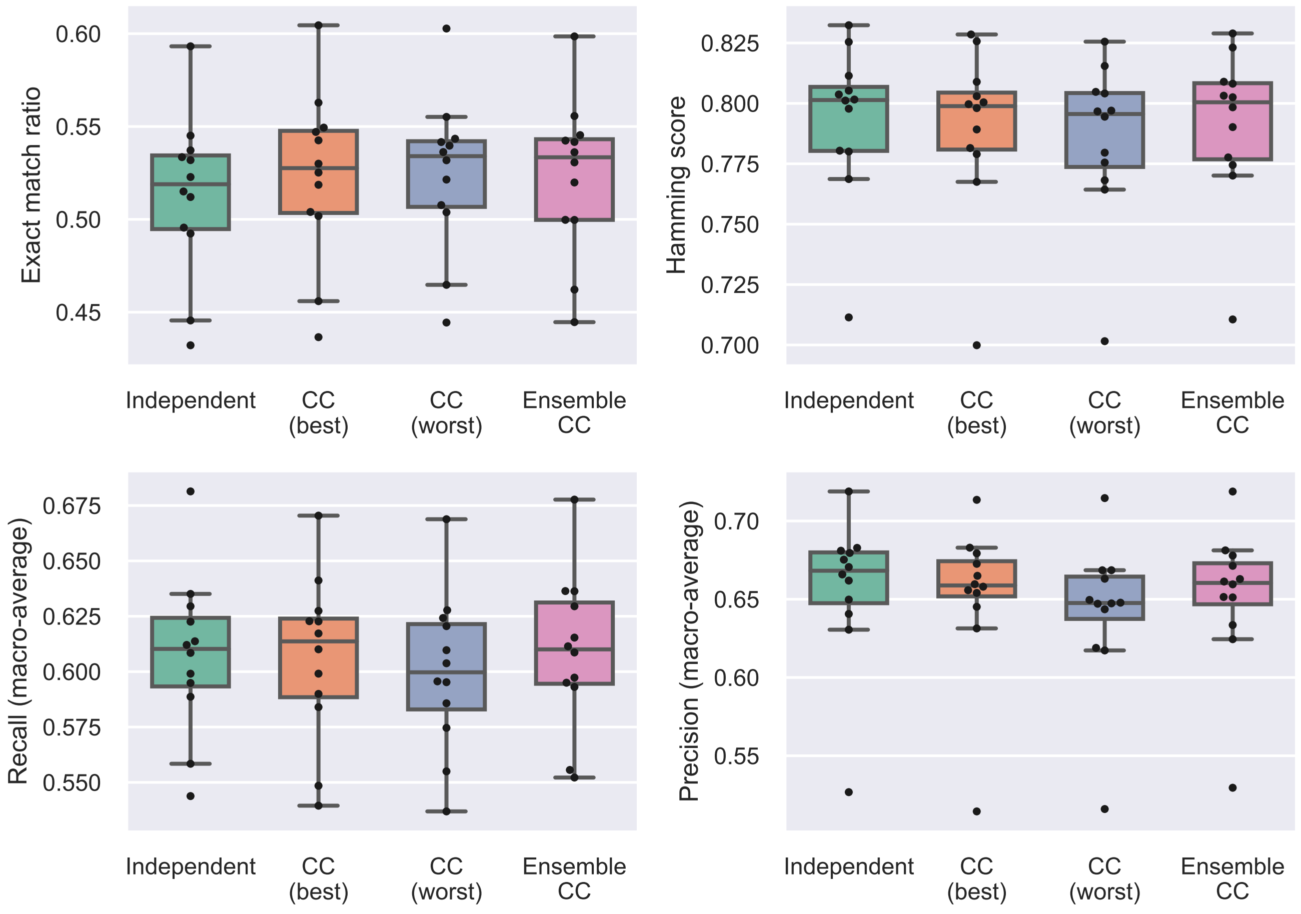


**Figure S1.** Classifier chain comparison using additional performance measures.

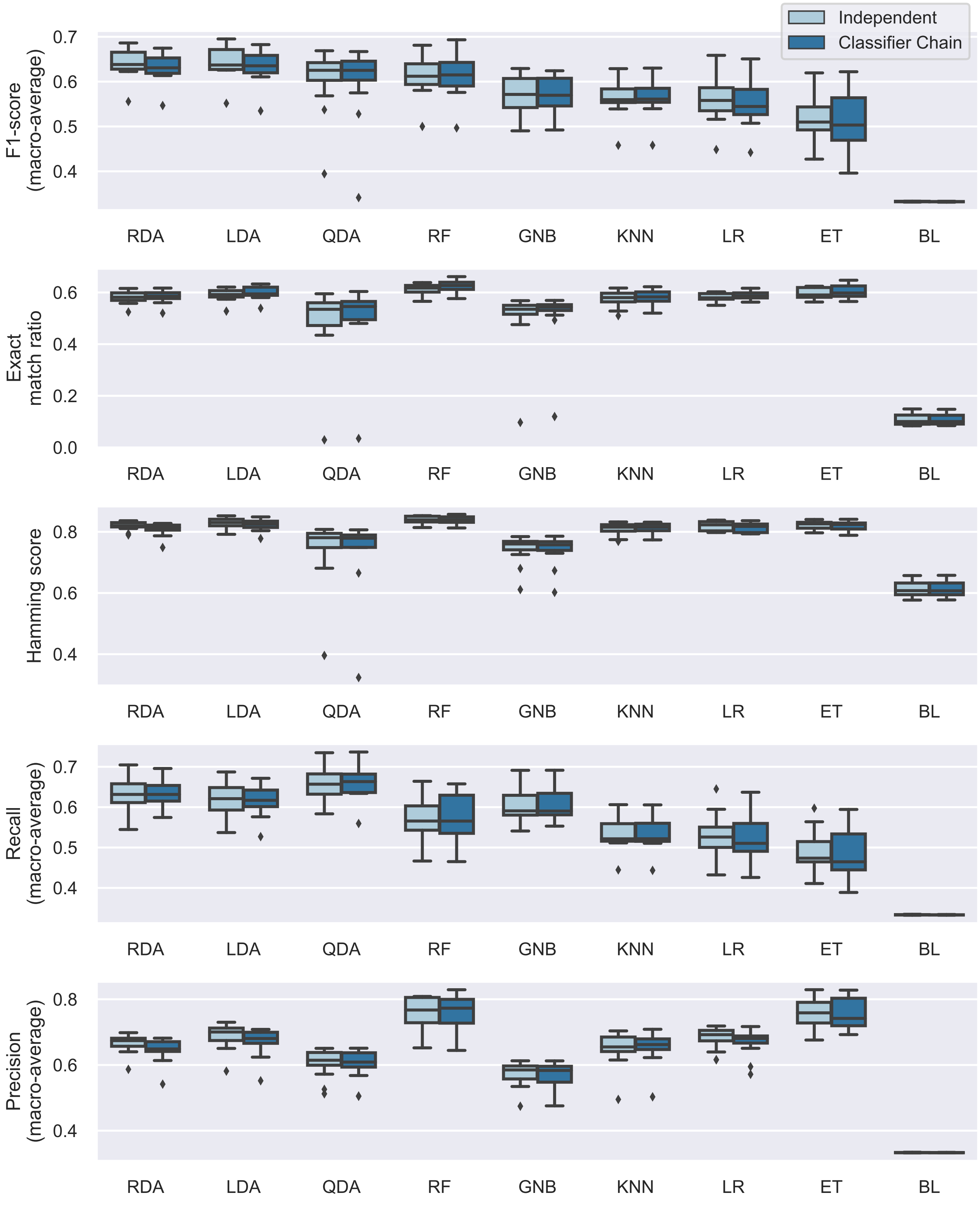


**Figure S2.** Multi-output independent classifier comparison using additional performance measures. Algorithms are presented in order of decreasing median macro-average F1-score.

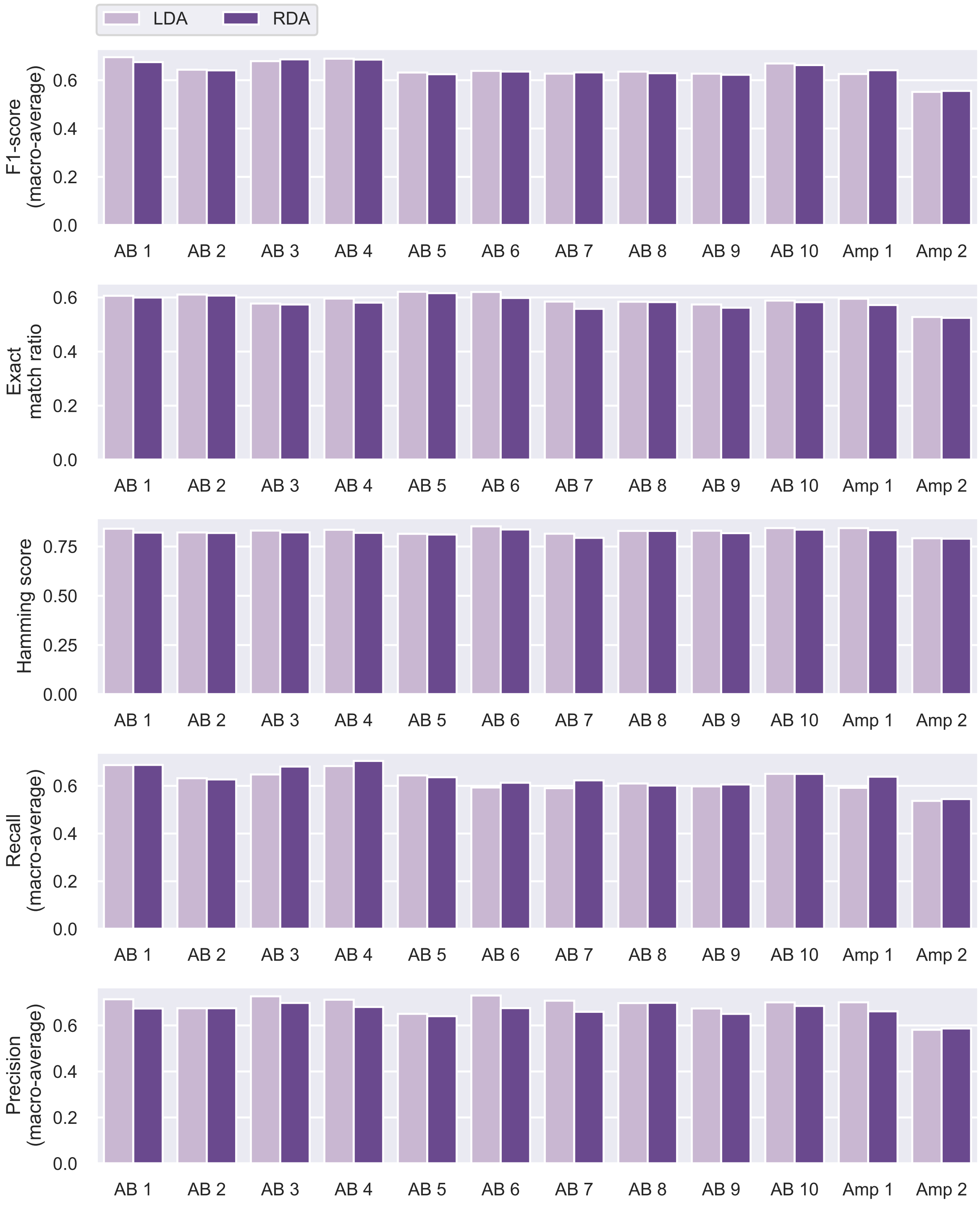


**Figure S3.** Multi-output independent classification performance with best-performing algorithms for individual participants. AB, able-bodied; Amp, amputee.

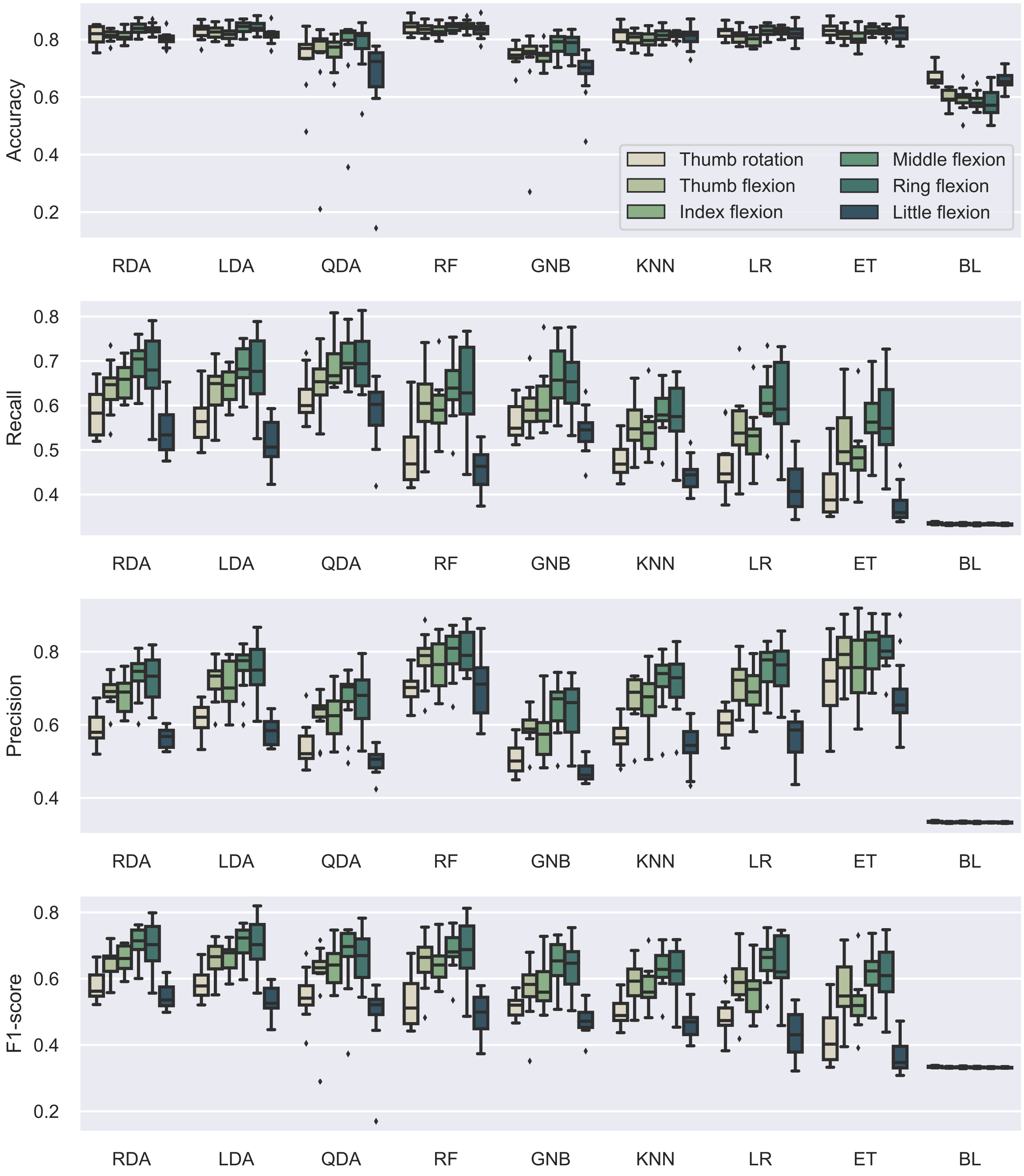


**Figure S4.** Multi-output independent classifier comparison for individual DOFs using additional performance measures. Algorithms are presented in order of decreasing median macro-average F1-score. Diamonds indicate outliers.

### Supplementary table

**Table S1.** Classification benchmark. Abbreviations, hyper-parameters, and search ranges.

| Classifier | Abbrv. | Hyper-parameter | Random search range |
|---|---|---|---|
| Baseline | BL | | |
| Logistic regression | LR | Regularisation parameter | logspace{min=$10^{-5}$, max=$10^{5}$, n=100} |
| Linear discriminant analysis | LDA | | |
| Quadratic discriminant analysis | QDA | | |
| Regularised discriminant analysis | RDA | Regularisation parameter $\alpha$ | linspace{min=0, max=1, n=101} |
| | | Regularisation parameter $\gamma$ | linspace{min=0, max=0.2, n=21} |
| Gaussian Naive Bayes | GNB | | |
| K-nearest neighbours | KNN | Number of neighbours | linspace{min=1, max=50, n=51} |
| | | Weights | {uniform, distance} |
| Random forests | RF | Number of estimators | {10, 20, 50, 100, 200, 500} |
| | | Maximum number of features | {auto, sqrt, log2} |
| | | Maximum depth | linspace{min=1, max=10, n=11} |
| | | Criterion | {gini, entropy} |
| Extra trees | ET | Number of estimators | {10, 20, 50, 100, 200, 500} |
| | | Maximum number of features | {auto, sqrt, log2} |
| | | Maximum depth | linspace{min=1, max=10, n=11} |
| | | Criterion | {gini, entropy} |
